# Frontal eye field inactivation alters the readout of superior colliculus activity for saccade generation in a task-dependent manner

**DOI:** 10.1101/646604

**Authors:** Tyler R. Peel, Suryadeep Dash, Stephen G. Lomber, Brian D. Corneil

## Abstract

Saccades require a spatiotemporal transformation of activity between the intermediate layers of the superior colliculus (iSC) and downstream brainstem burst generator. The dynamic linear ensemble-coding model (Goossens and Van Opstal, 2006) proposes that each iSC spike contributes a fixed mini-vector to saccade displacement. Although biologically-plausible, this model assumes cortical areas like the frontal eye fields (FEF) simply provide the saccadic goal to be executed by the iSC and brainstem burst generator. However, the FEF and iSC operate in unison during saccades, and a pathway from the FEF to the brainstem burst generator that bypasses the iSC exists. Here, we investigate the impact of large yet reversible inactivation of the FEF on iSC activity in the context of the model across four saccade tasks. We exploit the overlap of saccade vectors generated when the FEF is inactivated or not, comparing the number of iSC spikes for metrically-matched saccades. We found that the iSC emits fewer spikes for metrically-matched saccades during FEF inactivation. The decrease in spike count is task-dependent, with a greater decrease accompanying more cognitively-demanding saccades. Our results show that FEF integrity influences the readout of iSC activity in a task-dependent manner. We propose that the dynamic linear ensemble-coding model be modified so that FEF inactivation increases the gain of a readout parameter, effectively increasing the influence of a single iSC spike. We speculate that this modification could be instantiated by a direct pathway from the FEF to the omnipause region that modulates the excitability of the brainstem burst generator.

**Significance statement:** One of the enduring puzzles in the oculomotor system is how it achieves the spatiotemporal transformation, converting spatial activity within the intermediate layers of the superior colliculus (iSC) into a rate code within the brainstem burst generator. The spatiotemporal transformation has traditionally been viewed as the purview of the oculomotor brainstem. Here, within the context of testing a biologically-plausible model of the spatiotemporal transformation, we show that reversible inactivation of the frontal eye fields (FEF) decreases the number of spikes issued by the iSC for metrically-matched saccades, with greater decreases accompanying more cognitively-demanding tasks. These results show that signals from the FEF influence the spatiotemporal transformation.

## Introduction

The intermediate layers of the superior colliculus (iSC) are a key midbrain structure for generating saccadic eye movements, with the location of activity specifying the intended saccade vector (Gandhi and Katnani, 2011; Sparks, 2002; White and Munoz, 2011). Downstream of the iSC, this spatial coding is transformed into a temporal code, wherein the horizontal and vertical components of saccade displacement relate to the duration of recruitment of the brainstem burst generator. Goossens and Van Opstal (2006) proposed a dynamic linear ensemble-coding model that provides a biologically-plausible mechanism for this spatiotemporal transformation. In this model, each iSC spike contributes a fixed site-specific mini-vector to saccade displacement, so that the cumulative spike count monotonically increases along the intended saccade vector. In support of this model, the iSC emits an invariant number of spikes even during displacement-matched saccades that exhibit highly-perturbed kinematics due to an induced blink (Goossens and Van Opstal, 2006). This model also explains the non-linear main sequence relationship of peak saccade velocity to saccade amplitude via logarithmic coding of oculomotor space within the iSC and the associated rostral-caudal gradient of the temporal burst profiles of iSC neurons (Goossens and Van Opstal, 2012; Van Opstal and Goossens, 2008; Van der Willigen et al., 2011). More recently, Kasap and Van Opstal (2017, 2019) simulated the dynamics of iSC activity using a neural network model, and found that the inclusion of lateral synaptic interactions within the iSC produce plausible profiles of iSC activity once triggered by an external input, subsequently generating realistic saccades. This implies that once a saccade vector is specified, the iSC and downstream brainstem burst generator instantiate the spatiotemporal transformation for saccade generation.

The papers by Van Opstal and colleagues have considered the role of cortical and subcortical inputs into the iSC only from the perspective of specifying the intended saccade target. However, the iSC and other cortical (e.g., frontal eye fields; FEF, lateral intraparietal area) and subcortical (e.g., basal ganglia and cerebellum) areas have bidirectional connectivity and are also active during saccade generation (Barash et al., 1991; Bruce and Goldberg, 1985; Crapse and Sommer, 2009; Fuchs et al., 1993; Hikosaka et al., 2000; Wurtz et al., 2001). The FEF is particularly important, as its integrity is critical for saccade generation if the iSC has been permanently ablated or reversibly inactivated (Keating and Gooley, 1988; Schiller et al., 1980), presumably via FEF projections to the oculomotor brainstem that bypass the iSC (Segraves, 1992). These considerations have led us to wonder if a core prediction of the dynamic linear ensemble coding model, which is the generation of an invariant number of iSC spikes for displacement-matched saccades, would hold when the FEF is suddenly inactivated. In testing this prediction, we may gain insights into the potential role, or lack thereof, of the FEF in the spatiotemporal transformation for saccades.

In a recent series of studies, we have recorded iSC activity while reversibly inactivating the FEF with cryogenic cooling probes (Dash et al., 2018; Peel et al., 2017). This allowed us to examine the impact of a sudden decrease in FEF input while recording the same iSC neuron. Here, we test whether FEF inactivation alters the cumulative spike count in the iSC neurons for displacement-matched saccades. We do this across four different saccade tasks, since the impact of FEF inactivation increases for more cognitively-demanding tasks (Deng et al., 1986; Dias and Segraves, 1999; Peel et al., 2014; Sommer and Tehovnik, 1997). We find that FEF inactivation reduced the number of iSC spikes for displacement-matched saccades, with larger reductions accompanying more cognitively-demanding tasks. Importantly, the decrease in spike count extended throughout the saccade-related response field for a given iSC neuron. Fundamentally, the iSC emitted fewer spikes during FEF inactivation. These results demonstrate that signals from the FEF influence the readout of iSC activity by the brainstem burst generator, which we surmise relates to the excitability of the brainstem burst generator.

## Methods

This manuscript is partly based on experimental data reported in two previous studies (Dash et al., 2018; Peel et al., 2017), which characterized the effects of cryogenic FEF inactivation on iSC activity in immediate and delayed saccade tasks, respectively (Experiment 1). We also carried out novel experiments to address how FEF inactivation altered saccade-related response fields in the iSC (Experiment 2).

### Experimental procedures

Data was obtained from two male monkeys (*Macaca mulatta*, monkeys D, and O weighing 9.8, and 8.6 kg respectively) for each experiment. As previously described (Peel et al., 2017), each monkey underwent two surgeries to permit extracellular recordings from the iSC, and cryogenic inactivation of the FEF using cryoloops implanted into the arcuate sulcus. All training, surgical, and experimental procedures conformed to the policies of the Canadian Council on Animal Care and National Institutes of Health on the care and use of laboratory animals, and were approved by the Animal Use Subcommittee of the University of Western Ontario Council on Animal Care. We monitored the monkeys’ weights daily and their health was under the close supervision of the university veterinarians.

For each experiment, we recorded extracellular activity of an isolated iSC neuron pre-, peri-, and post-cooling of the FEF while monkeys performed a saccade task. Eye position signals were sampled using a single, chair-mounted eye tracker at 500 Hz (EyeLink II). All neurons were recorded ^~^1 mm or more below the surface of the SC, in locations where electrical stimulation (300 Hz, 100 ms, biphasic cathodal-first pulses with each phase 0.3 ms in duration) evoked saccades with currents < 50 μA. In the first experiment, potential cue locations were at the center of a neuron’s response field and at the diametrically opposition position. We approximated the center of a given neuron’s response field by identifying the saccade vector associated with the highest firing rates before FEF inactivation. To quantify the changes across the response field with FEF inactivation, we also conducted a second experiment wherein we recorded iSC neurons while monkeys performed saccades towards cues dispersed throughout the response field. We specified the number and spacing of cues (averages of 49 and 3°, respectively) within a square grid to sample the extent of a given neuron’s response field.

After completion of the pre-cooling session (^~^60 trials for each session), chilled methanol was pumped through the lumen of the cryoloops, decreasing the cryoloop temperature. Once the cryoloop temperature was stable at 3°C, we began the peri-cooling session. Cryoloop temperatures of 3°C silence post-synaptic activity in tissue up to 1.5 mm away without influencing axonal propagation of action potentials (Lomber et al., 1999). Upon finishing the peri-cooling session, we turned off the cooling pumps, which allowed the cryoloop temperature to rapidly return to normal. When the cryoloop temperature reached 35°C, we started the post-cooling session. Although saccadic behaviour and iSC activity rapidly recovered after rewarming, the post-cooling sessions may have contained residual effects of cooling. To minimize these and other time-dependent factors, we combined trials from pre- and post-cooling sessions into the *FEF warm* condition, which we compared to *FEF cool* condition (peri-cooling). We obtained similar results about the influence of FEF inactivation on iSC activity when only comparing the pre- and peri-cooling sessions.

### Behavioural tasks

We recorded iSC activity while monkeys performed either immediate (direct, or gap) or delayed (visually, or memory-guided) saccade tasks (n.b., we did not record any iSC neurons with both delayed and immediate saccade tasks). In all tasks, the monkeys first had to look at a central fixation point, and hold fixation within a ±3° window for a period of 750-1000 ms. On immediate saccade tasks, a peripheral cue was then presented either coincident with (direct saccade task) or 200 ms after (gap saccade task; central fixation was still required during this 200 ms gap) disappearance of the central fixation point. The monkeys then had 500 ms to look toward the peripheral cue within a spatial window (diameter set to 60% the visual eccentricity of the cue). The cue remained on for the direct saccade task, but was flashed for 150 ms in the gap saccade task. In the delayed saccade tasks, the central fixation point remained on following peripheral cue presentation for a fixed delay period of 1000 ms, during which the monkeys maintained central fixation. Peripheral cues were either extinguished after 250 ms (memory saccade task) or remained on for the remainder of the trial (delayed visually-guided saccade task). Monkeys had to saccade to the remembered (memory saccade task) or visible cue (delayed visually-guided saccade task) location (window diameter set to 70% the visual eccentricity of the cue) within 1000 ms after offset of the central fixation point, which served as the go-cue. Saccade onset and offset were determined using a velocity criterion of 30°/s. We excluded trials from our analysis where monkeys did not generate their first saccade towards the target, generated anticipatory saccades with saccadic reaction times < 60 ms, or blinked during the trial.

For Experiment 1 where we characterized the effects of FEF inactivation on iSC activity at the center of neuron’s response field, we usually recorded neurons with only one type of delayed saccade (i.e., ^~^79% of iSC neurons were recorded with either memory or delayed visually-guided saccades), but we always recorded iSC neurons with both direct and gap saccades in an interleaved manner. Furthermore, we only recorded neurons with a single saccade task for Experiment 2 that sought to characterize the effects of FEF inactivation on neuronal response fields. Of the 29 isolated neurons that were studied in Experiment 2 and exhibited a clear response field, we recorded 8, 12, and 9 neurons with direct, delayed visually-guided, and memory-guided saccades respectively. Given these small sample sizes and that FEF inactivation produced similar effects across tasks, we pooled results across saccade task in Experiment 2.

### Neuron classification

In this study, we analyzed neurons exhibiting saccade-related activity. For classification purposes, we convolved individual spikes with a spike density function that mimics an excitatory post-synaptic potential (rise-time of 1 ms, decay-time of 20 ms, kernel window of 100 ms (Thompson et al., 1996)). To qualify as a saccade-related neuron, the mean peri-saccadic firing rates (defined in an interval spanning 8 ms before saccade onset to 8 ms prior to its end) had to be significantly greater than the last 100 ms before the go-cue (p < 0.05, Wilcoxon signed rank test), and the increase in peri-saccadic activity above baseline activity (last 200 ms before fixation cue offset in immediate response tasks, or last 200 ms before cue onset in delayed response tasks) had to exceed 50 spikes/s (McPeek and Keller, 2002; Munoz and Wurtz, 1995; Peel et al., 2017). Because the delayed saccade tasks allowed us to dissociate visual and saccade-related activity, we also differentiated visuomotor from motor neurons by looking for the presence or absence, respectively, of significantly increased visually-related activity (mean activity in a 50 ms window after the onset of any visually-related activity) compared to the last 200 ms before peripheral cue onset (p < 0.05, Wilcoxon signed rank test, see (Peel et al., 2017) for more details on quantification of visually-related activity).

### Dynamic linear ensemble-coding model

To explore the impact of FEF inactivation on iSC activity within the context of the dynamic linear ensemble-coding model (Goossens and Van Opstal, 2006), we compared the cumulative number of saccade-related spikes for saccades of similar metrics within the four tasks we studied, with or without FEF inactivation. One prediction from the dynamic linear ensemble-coding model is that metrically-matched saccades should be associated with an equivalent number of spikes across the population of iSC neurons. Within a single neuron, we tallied the cumulative number of saccade-related spikes within a window spanning 20 ms before saccade onset to 20 ms before saccade offset (similar results were found if we used an interval shifted 8 ms relative to saccade onset and offset). The dynamic linear ensemble-coding model follows the framework of Equation 1 whereby each spike (*k* = 1, 2, …, *N*) from each iSC neuron (*i* = 1, 2, …, *M*) adds a fixed site-specific amount 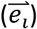 convolved with a delta function (*δ*) to the saccade vector following a delay:

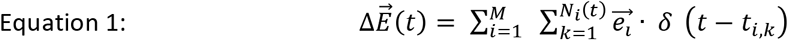

In an implementation of the model described previously (see Figure 6 of Goossens and Van Opstal, 2006), the brainstem circuits below the iSC simply read out spatial activity for horizontal and vertical components in two independent linear feedback systems; normal saccade dynamics can be realized after optimizing five fixed parameters (scaling factors for each horizontal and vertical component based on projection strength, feedforward gain of saccade burst neurons, and fixed delays of brainstem activation and of the feedback loops). Importantly, more recent studies implementing this model (Van der Willigen et al., 2011) have found that only two free parameters, the feedforward gain of the burst generator and the delay of brainstem activation, were sufficient to create realistic saccades.

### Matched-saccade procedure

FEF inactivation increases the RT and decreases the accuracy and velocity of contralateral saccades (Peel et al., 2014). Even with such changes, the distributions of these parameters with or without FEF inactivation overlap, and we exploit this overlap in the current study. As described below, one of the means by which we test the model across FEF inactivation is to find instances where a very similar saccade is generated with or without FEF inactivation, while the same neuron was being recorded. To compare cumulative spike counts of such metrically-matched saccades, we first ranked each FEF cool trial with FEF warm trials based on the difference in horizontal and vertical displacement. We then matched with replacement each FEF cool trial with the FEF warm trial that contained the lowest combined rankings (we used this matching procedure in our previous study (Peel et al., 2017), modifying it here to not include a match of saccade peak velocity). We specified that any such matched saccades had to have differences of horizontal and vertical displacements less than 1.5°, but matches usually had differences in horizontal and vertical displacements much less than 1° (mean ± SD of −0.001 ± 0.322° and −0.001 ± 0.317° across 4,460 matched pairs of trials with ipsilesional iSC recordings, respectively). In some analyses, we also tested whether this matching procedure biased any results by performing a similar matching procedure utilizing only FEF warm trials. To do this, we matched each FEF warm trial to a different FEF warm trial that had the lowest ranked differences in horizontal and vertical saccade displacements.

### Characterization of iSC response fields

The dynamic linear ensemble-coding model posits each saccade-related spike from the population of active iSC neurons adds a fixed, site-specific displacement vector to the saccade trajectory; in doing so, the model makes no assumptions about the spatial distribution of saccade-related activity throughout the iSC. Hence, it is possible that the effects of FEF inactivation on spike counts at the center of a given neuron’s response field (which we test in Experiment 1) could be confounded by coincident changes to the tuning width of iSC response fields (i.e., expansion, or shrinking). From the additional 29 isolated iSC neurons recorded for Experiment 2, we examined the influence of FEF inactivation on saccade-related response fields. To do this, we first constructed separate response fields for neural activity recorded during saccades generated with or without FEF inactivation. We then found the average spike count for each point ± 2° within a 2D grid of saccade displacement (spacing of 1°), normalizing data based on the FEF warm condition (subtracting minimum value and then dividing by max value), and then linearly interpolating data to create a 2D colourmap. We then identified various contour levels of spike counts (0.3, 0.5, 0.7, and 0.9; 1.0 represents peak spike count from the FEF warm condition) in each of the FEF warm and cool conditions, which provided a comparative measure for how FEF inactivation influenced the center and tuning width of a neuron’s response field.

To better quantify changes in spike counts during FEF inactivation, we also employed a static response field model of Ottes et al. (Ottes et al., 1986), which is illustrated by Equation 2.

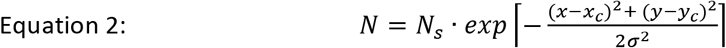

Parameter *N_s_* represents the maximal spike count in the response field, which is located at *x_c_* and *y_c_* within the iSC map (center position defined in mm). The decay rate or tuning width of the response field is governed by *σ* (also defined in mm and corresponding to the radial distance from the response field center to ^~^61% of peak response). For each neuron and cooling condition, we identified the values of four parameters (*N_s_*, *x_c_*, *y*_c_, and *σ*) that satisfied the least squares criterion using a nonlinear optimization algorithm (Goossens and Van Opstal, 2006). Of the 29 saccade-related response fields tested in this manner, we only excluded 3 iSC neurons for further analyses that did not produce reasonable estimates of the maximal spike count parameter (*N_s_* between 5 and 50 spikes in the FEF warm condition). Importantly, this static field model of response fields obtained robust fits to FEF warm data (Pearson’s correlation ranged between 0.55 and 0.87 across our sample of 26 neurons), which did not significantly change with FEF inactivation (p = 0.62, z = −0.4978, Wilcoxon signed rank test). Average parameter values ± standard deviation for *N_s_*, *x_c_*, *y_c_*, and *σ* in the FEF warm condition were 12 ± 6 spikes, 2.7 ± 1.2 mm, 0.2 ± 0.8 mm, and 0.9 ± 0.3, respectively. While our average spike counts and tuning widths are lower compared to values (*N_s_* = 20 ± 9 spikes, *σ* = 0.5 ± 0.2 mm) reported by (Goossens and Van Opstal, 2006), such differences may relate to the reduced vigor of iSC activity in the delayed saccade tasks compared to immediate saccade tasks employed in the earlier study by Goossens and Van Opstal (2006). Finally, because changes in these parameters may covary with FEF inactivation, we also tested whether FEF inactivation induced any expansion or shrinkage of response field width (*σ*). To test this, we performed a second fitting procedure on the FEF cool data using fixed parameters of *N_s_*, *x*_c_, and *y_c_* extracted from the FEF warm fit, and only allowed the remaining *σ* parameter to vary.

### Experimental design and statistical analysis

Our analysis of the effects of FEF inactivation on iSC activity exploits the overlap of saccade vectors with and without FEF inactivation, such that we could examine differences in iSC activity between matched trials having similar saccade vectors towards the center of the response field. To quantify the effects of FEF inactivation on saccadic behaviour and related measures of iSC activity, we usually performed paired Wilcoxon singed-rank tests to find statistical differences within individual neurons and across the neuronal population at p < 0.05. We also investigated how FEF inactivation influenced the response fields of iSC neurons by comparing parameters generated from a response field model (Ottes et al., 1986) with and without FEF inactivation. We used paired Wilcoxon singed-rank tests to uncover statistical differences across individual iSC neurons and across the neuronal population at p < 0.05.

### Results

In this study, we tested if the dynamic linear ensemble-coding model proposed by (Goossens and Van Opstal, 2006) is robust to FEF inactivation across four saccade tasks (direct, gap, delayed visually-, and memory-guided saccades). In experiment 1, we focus on activity recorded from 150 saccade-related iSC neurons (83 from Monkey DZ, and 67 from Monkey OZ) recorded ipsilateral to the side of FEF inactivation while monkeys performed saccades towards peripheral cues placed at the center of a neuron’s response field. Although FEF inactivation produced a triad of saccadic defects for contralaterally-directed saccades (reduced accuracy, and peak velocities, and increased reaction times) in every task, a substantial overlap of saccade metrics allowed us to test the model for saccades matched closely for saccade metrics. Indeed, our matching procedure matched 96% of FEF cool trials with a FEF warm trial containing very similar saccade metrics (see **Methods** for details). We first demonstrate how FEF inactivation influenced spike counts in an exemplar iSC neuron for matched saccades in a single task, then we characterize the effects of FEF inactivation across the sample of recorded iSC neurons, and across different saccade tasks. In experiment 2, using data from an additional 29 iSC neurons recorded ipsilateral to FEF inactivation, we examined the impact of FEF inactivation on neurons’ movement fields within the context of the model.

### FEF inactivation reduced cumulative spike counts in iSC neurons for displacement-matched saccades

In contrast to what would have been predicted by the model if it were to operate independently of any supra-tectal inputs, FEF inactivation reduced the cumulative spike counts for saccades of matched displacement. **Figure 1A** shows data for a pair of matched saccades from the delayed visually-guided saccade paradigm. These saccades were closely matched for horizontal and vertical displacement (displacement differences less than 0.1° for FEF warm and cool conditions). As expected, FEF inactivation decreased peak saccade velocity (from 709 to 527 °/s in this case) and increased saccade duration (from 49 to 53 ms). Despite the similarity in saccade metrics, FEF inactivation markedly reduced the cumulative spike count from 24 to 12 spikes during the interval spanning from 20 ms before saccade onset to 20 ms before saccade offset (see shaded area with spike trains in **Figure 1A**).

**Figure 1.**
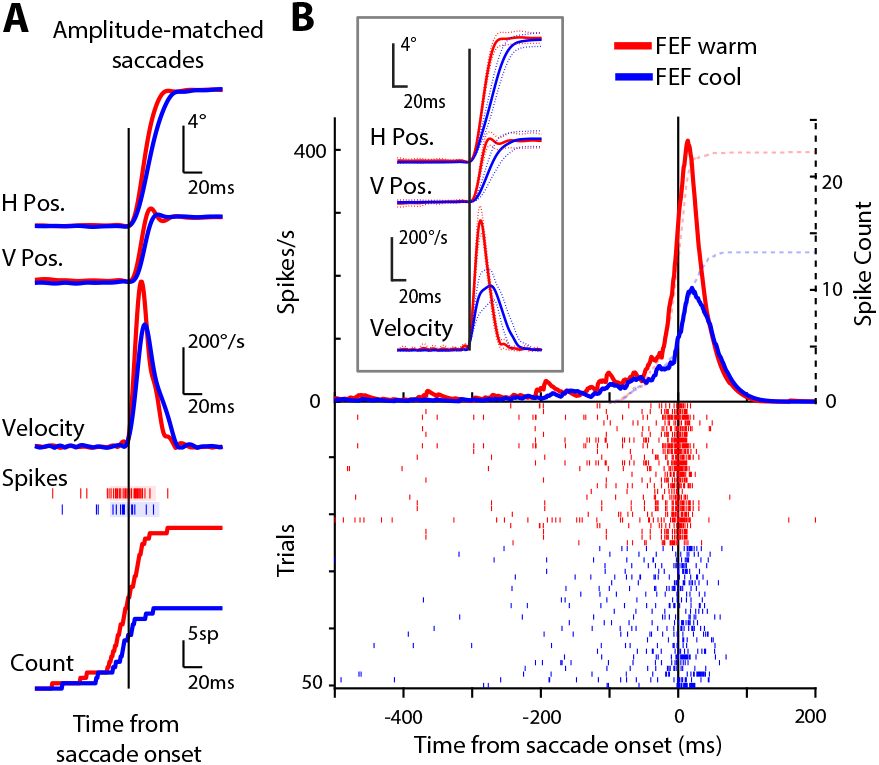
FEF inactivation reduces the cumulative spike counts in iSC neurons for metrically-matched saccades. **(A)** Spike rasters and cumulative density functions (CDF) from one matched pair of delayed visually-guided saccades. Our matched saccade analysis compared saccades of very similar eye position profiles but different kinematics and reaction times across FEF warm or FEF cool conditions. Shaded region within spike train indicates period between saccade onset to offset. **(B)** Spike rasters (below) and mean spike density functions (above) showing reduced cumulative spike counts in an example ipsilesional iSC neuron during FEF inactivation. Note how metrically-matched, delayed visually-guided saccades had reduced peak velocities, and longer durations with FEF inactivation (inset). Spike count indicates the cumulative number of spikes starting 100 ms before saccade onset.

We repeated this procedure for all trials recorded from this neuron, matching saccades generated in the FEF cool condition to one generated in the FEF warm condition; doing so yielded 25 matched pairs. The mean (+/- SE) displacement and radial velocity of these matched pairs is shown in the inset of **Figure 1B**, emphasizing the close match in saccade displacement despite decreases in saccade velocity and corresponding increases in saccade duration. Across all matched trials recorded from this iSC neuron (plotted in order in the rasters of **Figure 1B**), we observed a significant decrease in the cumulative number of spikes from 17 to 10 in the FEF warm to cool conditions (p < 0.0001, z = 3.9594, Wilcoxon signed-rank test), despite a significant increase in saccade duration from 48 to 65 ms (p < 0.0001, z = −4.3759, Wilcoxon signed-rank test). Hence, despite increases in saccade duration and the generation of metrically-matched saccades, the total number of iSC spikes decreased during FEF inactivation.

### Cumulative spike counts for metrically-matched saccades decreased during FEF inactivation in a task-dependent manner

The neuron shown in **Figure 1B** exhibited one of the larger changes in cumulative spike count during FEF inactivation. Across the sample of recorded neurons, FEF inactivation reduced cumulative spike count during the saccade interval, with larger reductions in spike count accompanying saccades generated during more cognitively demanding saccade tasks. Of the four saccade tasks, FEF inactivation caused the largest reductions in the cumulative spike count during memory-guided saccades (black dots in **Figure 2A**; cumulative spike count decreased by 10% across 58 iSC neurons; p < 0.0001, z = 4.0522, Wilcoxon signed-rank test), and the next largest reductions during delayed visually-guided saccades (grey dots in **Figure 2A**; cumulative spike count decreased by 5% across 58 iSC neurons; p < 0.01, z = 2.8909, Wilcoxon signed-rank test). Since these tasks dissociate visual and saccade-related activity, we investigated whether the reduction in cumulative spike count during FEF inactivation related to the cell’s functional classification. Although FEF inactivation produced larger decreases of cumulative spike counts in pure motor neurons compared to visuomotor neurons, these spike count differences between neuron classifications did not significantly differ in the memory- (p = 0.33, z = −0.9708, Wilcoxon rank sum test) or delayed visually-guided saccade tasks (p = 0.85, z = −0.1866, Wilcoxon rank sum test).

**Figure 2.**
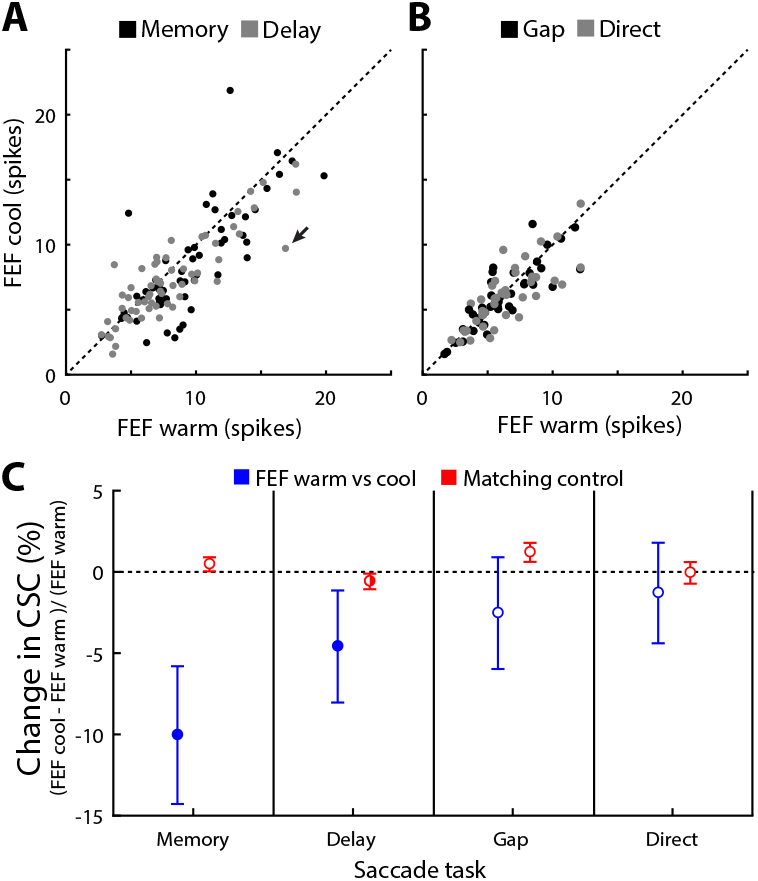
The reduction in iSC spike count during FEF inactivation scaled with task demands. **(A)** FEF inactivation consistently decreased the cumulative number of spikes during saccades (20 ms before saccade onset to 20 ms before saccade offset) across the population of ipsilesional iSC neurons for both memory- and delayed visually-guided saccades. **(B)** FEF inactivation often reduced the cumulative spike count during gap and direct saccades, but these decreases were not consistent across the population of iSC neurons. **(C)** FEF inactivation caused greater and more consistent decreases of cumulative spike count depending upon the saccade task (blue lines, mean ± SE). Such inactivation effects were most commonly observed for memory saccades followed by delayed visually-guided, gap, and finally, direct saccades, but did not occur with a similar matching procedure of only FEF warm trials (red lines). Filled and half-filled circles indicate significant results at 0.01 and 0.05 criterions, respectively.

Compared to the results in the delayed saccade tasks, FEF inactivation decreased cumulative spike counts in the gap and direct saccade tasks by a smaller amount of 3 and 1%, respectively, and these effects did not reach significance across our sample (**Figure 2B**, both 43 neurons, p = 0.39 and 0.17, z = −0.8556 and −1.3880, respectively, Wilcoxon signed-rank tests).

The impact of task is summarized in **Figure 2C** (blue bars), emphasizing how FEF inactivation produced larger and more consistent reductions of cumulative spike count for saccade tasks with increasing cognitive demands (i.e. delaying response, and remembering peripheral cue location). As a control, we repeated our matching procedure using data only from FEF warm trials (i.e., a given FEF warm trial would be paired with a different FEF warm trial of closely matched displacements); doing so reveals the amount of noise inherent to the matching procedure. Across all four saccade tasks, cumulative spike counts did not change (all differences less than 1.2%, p > 0.01, Wilcoxon signed-rank tests) when FEF warm trials were matched to other FEF warm trials (red bars and symbols, **Figure 2C**).

The above analysis averages the number of saccade-related spikes within a given neuron for qualifying matched saccades, and then plots the change across FEF inactivation on a neuron-by-neuron basis. We also performed an analysis where each matched pair was treated as its own sample; doing so gives a sense of the trial-by-trial variability inherent to FEF inactivation, and to the matching procedure. The results of this analysis are shown in **Figure 3A** for the memory-guided saccade task. Here, the size of each square is proportional to the number of trials with the observed number of spikes across matched pairs; for example, the square indicated by the grey arrow shows 5 matched trials (across all of our sample) that had cumulative spike counts of 14 and 8 spikes in the FEF warm and FEF cool condition, respectively. While this analysis shows considerable variation around the line of unity, FEF inactivation shifted the cumulative spike counts towards reduced values (blue histograms, 10% decrease, p < 10^−17^, z = 8.6837, matched pairs = 1151, Wilcoxon signed-rank test). In contrast, we observed negligible shifts when we matched only FEF warm trials in the memory-guided saccade task (red histograms in **Figure 3A**, p = 0.93, z = −0.093781, matched pairs = 2643, Wilcoxon signed-rank test).

**Figure 3.**
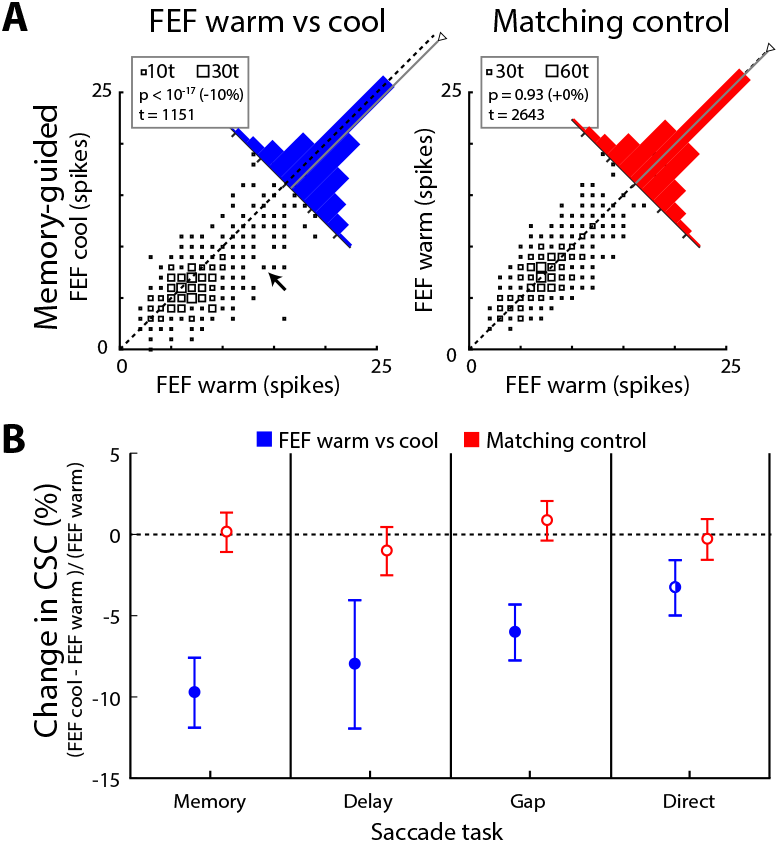
A trial-by-trial analysis of iSC spike counts revealed reductions during FEF inactivation across all tasks. **(A)** As shown for the memory-guided saccade task, FEF inactivation reduced cumulative spike counts across trials matched for saccade metrics (blue bars). Note how histograms are skewed towards decreased cumulative spike counts. In contrast, we found invariant cumulative spikes when matching only FEF warm trials (red bars). Each square represents the occurrence of a given pairing of spike counts, with number of occurrences of a particular pairing represented as the size of the square (see insert for reference). **(B)** Across trials, FEF inactivation produced larger relative reductions in cumulative spike count depending upon saccade task (mean ± SE). Same format as Fig. 2C.

When analysed this way, we observed significant reductions in cumulative spike counts in all four tasks when FEF warm and FEF cool trials were matched (blue symbols and lines in **Figure 3B**), but not when FEF warm trials were matched to FEF warm trials (red symbols and lines in **Figure 3B**). As before, the magnitude of reduction varied with the saccade task, with the largest decreases accompanying saccades generated in the memory-guided saccade task (−10%, p < 10^−17^, z = 8.6837, Wilcoxon signed-rank test), then the delay task (−8%, p < 10^−2^, z = 2.8909, Wilcoxon signed-rank test), then the gap task (−6%, p < 10^−2^, z = −2.8553, Wilcoxon signed-rank test), and finally the direct-saccade task (−3, p < 0.05, z = −2.0153, Wilcoxon signed-rank test). Recall that the cue was flashed for only 150 ms for gap but not direct saccades, hence saccades in the gap saccade task did not always land on a visible target (^~^54% of gap saccades had RTs greater than 150 ms without FEF inactivation).

Collectively, the results in **Figures 2** and **3** show that FEF inactivation decreased the cumulative number of spikes in ipsilesional iSC neurons for displacement-matched saccades in a task dependent manner. Next, we evaluated whether spike-count reductions in the ipsilesional iSC during FEF inactivation could be related to changes in fixation position.

### Differences in fixation position cannot explain cumulative spike count decreases during FEF inactivation

FEF inactivation produces slight deviation in fixation position towards the intact side (see Figure S1 of Peel et al., 2016). Could the reduction in spike count for displacement-matched saccades be related to such changes in fixation position? To explore this question, we divided matched pairs of trials into two subsets using a median-split procedure of fixation error when the FEF was inactivated (see **Figure 4A** for the segregation of high and low fixation errors at saccade onset for FEF cool trials). Doing so created one subject of FEF warm trials matched to FEF cool trials with a larger-than-average fixation error, and another subset of FEF warm trials matched to FEF cool trials with a smaller-than-average fixation error. As expected, the fixation error for the higher-than-average subgroup was significantly greater during FEF cool versus FEF warm trials (increase of 0.5°, p < 10^−38^, z = 13.0688, Wilcoxon sign-rank test). Critically, the fixation error for the lower-than-average subset was significantly *less* during FEF cool versus FEF warm trials (decrease of 0.2°, p < 10^−11^, z = −6.8679, Wilcoxon sign-rank test). We then analyzed the reductions in spike count for these two subsets, and found that cumulative spike count in the SC decreased to the same degree during FEF inactivation, regardless of the magnitude of any fixation error (**Figure 4B** and **C** show that spike count decreased by 9 or 11% for the greater-than-average or lower-than-average FEF cool fixation error in the memory-guided saccade task, respectively; p < 10^−7^ Wilcoxon signed-rank test for both subgroups, z = 5.3474 and 6.8254, respectively). We found a similar lack of effect of fixation error for all saccade types, and for different measures of fixation error (e.g., averaged during the entire pre-cue period, or when the horizontal or vertical component of fixation error was analyzed separately; data not shown). Overall, these analyses emphasize that changes in fixation error cannot explain the reductions in cumulative spike count in the SC during FEF inactivation.

**Figure 4.**
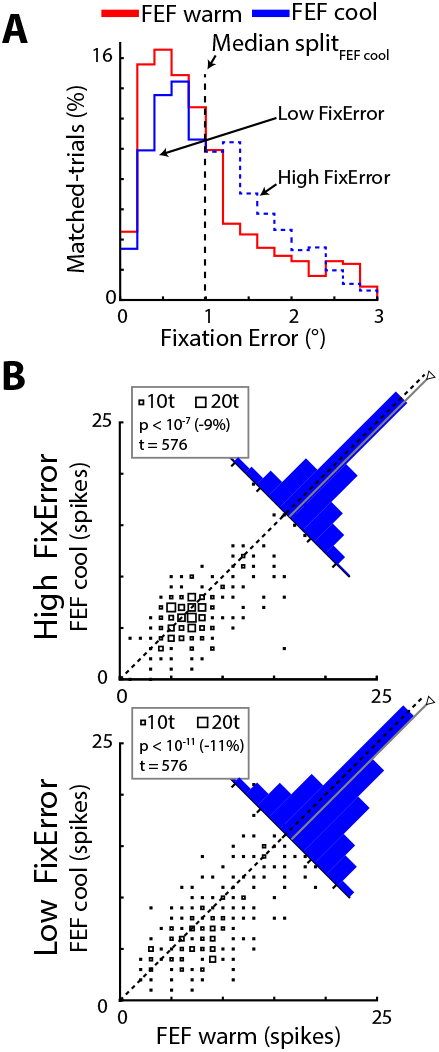
FEF inactivation reduces iSC spike counts for metrically-matched saccades regardless of fixation position. **(A)** As shown across all matched trials in the memory-guided saccade task, FEF inactivation (trials represented in blue) skewed the distribution of radial position error at the start of the saccade towards larger errors (compare to FEF warm trials represented in red). The blue vertical line represents the median value from FEF cool trials, which we used for the median-split analysis. **(B)** For both subsets of matched trials containing either high (top) or low (bottom) fixation error values in this task, FEF inactivation reduced cumulative spike counts as shown by the rightward skewness in the histograms. Same format as **Fig. 3A**, separated into subsets with higher- or lower-than average fixation error.

### FEF inactivation reduces the cumulative number of saccade-related spikes throughout the response field

Up to now, our analyses have focused on matched saccades generated toward cues placed at the estimated center of a neuron’s response field. However, the dynamic linear ensemble-coding model of Goossens and Van Opstal (2006) is based on the population of spike activity across all iSC neurons active for a given saccade. Theoretically, the decreases in spike count from the center of the response field could be offset by increases in spikes by shifts or expansion of the response field. To explore these possibilities, we analyzed data in experiment 2 where we characterized the entire saccade-related response field with and without FEF inactivation.

We constructed saccade-related response fields (see **Methods** for more details) with and without FEF activation, normalizing all data to the peak of the FEF warm response field for analysis across our sample. We then compared a number of parameters of the response field across FEF inactivation. An example of the saccade-related response fields from a representative neuron recorded with or without FEF inactivation is shown in **Figure 5A**. In this example, and consistent with our previous results, we found that FEF inactivation decreased the peak cumulative spike count during the saccadic interval for movements to the center of the response field (from 9 to 7 spikes), and other measures of peak saccade-related activity (in this case, 181 to 125 spikes/s). Moreover, the various contour levels (0.3 to 0.9 in 0.1 increments) reveals that FEF inactivation did not drastically alter the shape or extent of the movement field of this iSC neuron.

**Figure 5.**
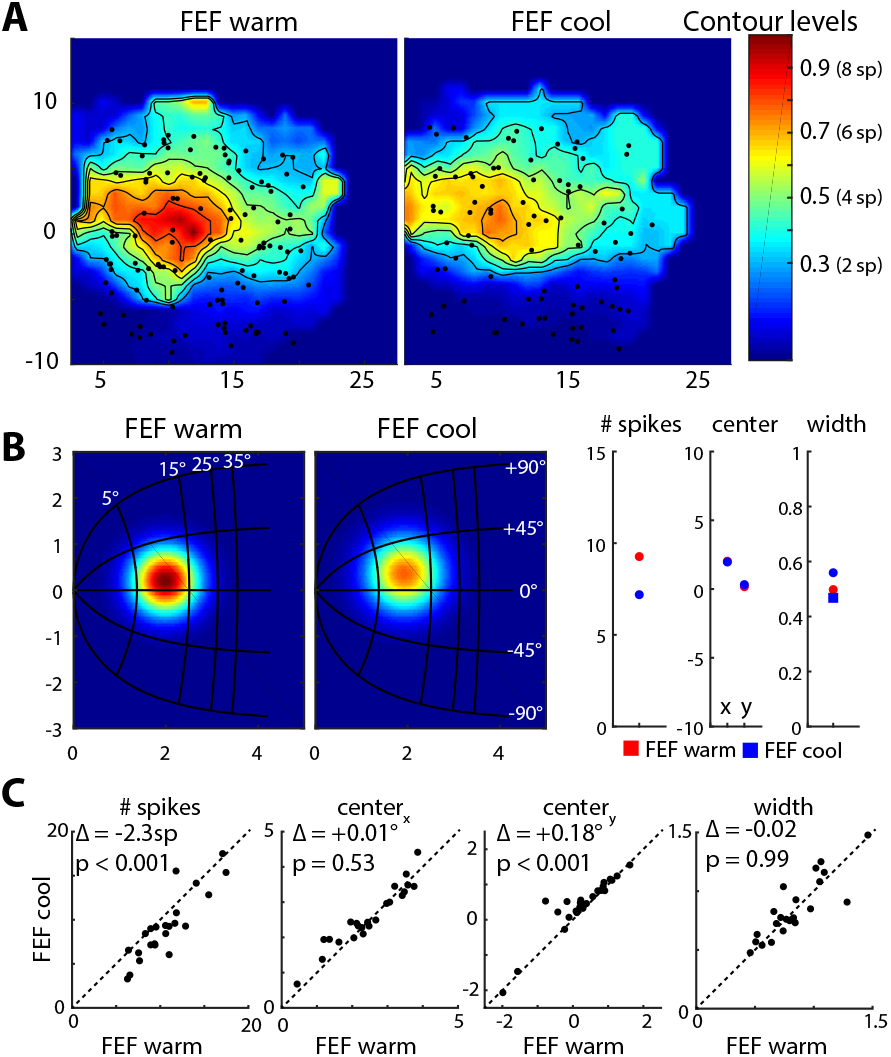
FEF inactivation reduces the overall spike count of iSC neuron population. **(A)** As shown for an example iSC neuron, FEF inactivation decreased spike count across its response field. As further detailed in the **Methods**, we characterized response fields two ways. Firstly, we performed a linear interpolation procedure of spike counts for each measured saccade displacement, where data within each plot is normalized to the maximum spike count in the FEF warm condition (value of 1). Shrinking of fixed contour lines with FEF inactivation reveal a blunting of spike counts at the center of the neuron’s response field. **(B)** Secondly, we employed a static response field model to quantify changes in this example neuron, whereby we used a least squares criterion within a nonlinear optimization algorithm to search for four key parameters (peak spike count, horizontal and vertical position in SC map, and tuning width) characterizing response fields in each cooling condition. Note that we used a standard representation of visual space within the iSC as implemented within the model, but this is likely an over-simplification for the upper and lower visual field (Hafed and Chen, 2016). In this example neuron, FEF inactivation reduced the peak spike count with only modest differences in other model parameters. An additional fit of FEF cool data using parameter values extracted from fits to the FEF warm condition except for tuning width revealed a negligible influence of FEF inactivation on tuning width (square symbol). This is consistent with the modest changes in tuning width during FEF inactivation being due to coincident changes in other parameters, including the blunting of the peak spike count. **(C)** Across our sample of 26 iSC neurons, FEF inactivation consistently reduced the peak spike count across our sample, with modest effects to the vertical position of response fields within the SC map. Importantly, the tuning widths of response fields were largely unaffected during FEF inactivation.

To quantify these effects across our sample, we employed a nonlinear optimization algorithm on a static response field model of Ottes et al. (1986) to obtain parameters that characterized key aspects of the response field (peak spike count, *N_s_;* tuning width, *σ;* horizontal and vertical center position in SC, *x_c_* and *y*_c_, respectively, see **Methods** for details). The results from this analysis on the same neuron are shown in **Figure 5B** which again shows how FEF inactivation blunted the peak spike count (9 to 7 spikes) at the response field center without affecting this position within the SC map (horizontal and vertical differences less than 0.2 mm). While FEF inactivation slightly increased the tuning width from 0.50 to 0.56 mm, such increases may be due to coincident changes in the other three parameters. Specifically, in a second fit of FEF cool data where we fixed the *N_s_*, *x_c_* and *y_c_* parameters based on FEF warm data (hence only *σ* could vary), we in fact found a smaller tuning width during FEF inactivation (0.47 mm). This illustrates how FEF inactivation primarily blunted the peak spike count of iSC response fields without systematically changing its tuning width, suggesting that the overall spike count from this neuron decreased regardless of saccade vector.

Across 26 of 29 iSC neurons that had reasonable estimates of peak spike count in the FEF warm data (*N_s_* between 5 and 50 spikes), we found similar changes to parameters with FEF inactivation (**Figure 5C**). FEF inactivation significantly reduced the peak spike count across our sample from 12 to 10 (p < 0.001, z = 3.3499, Wilcoxon signed-rank test), but had no overall effect on tuning widths (0.88 to 0.86 mm, p = 0.99, z = −0.0135, Wilcoxon signed-rank test). Likewise, after performing an additional fit on FEF cool data using fixed parameters except for tuning width, we again found that tuning widths remained unchanged during FEF inactivation, although there was a trend toward decreasing widths of response fields (p = 0.058, z = 1.8969, Wilcoxon signed-rank test). Together, the effects of FEF inactivation on the response field parameters of peak spike count and tuning width suggest that the overall spike count decreases throughout the entirety of the response field, regardless of saccade vector. While the horizontal position for the response field center did not change during FEF inactivation (p = 0.53, z = - 0.6323, Wilcoxon signed-rank test), we did observe a significant upward bias for its vertical position (p < 0.001, z = −3.5652, Wilcoxon signed-rank test), although the magnitude of this change was quite small (0.2 mm, or 2.5 times smaller than the average tuning width of iSC neuron with a value of 0.5 mm). Given that FEF inactivation had no impact on the tuning widths of iSC response fields, this would seem to preclude the possibility that decreases in the cumulative spike count for metrically-matched saccades could be explained by additional spikes from the margins of the response field. Likewise, while modest vertical biases in iSC response fields during FEF inactivation was an unexpecting finding, such a mechanism by itself would not increase the overall spike count from the population of iSC neurons, as additional spikes for saccades toward the upper visual field would be offset by reductions for downward saccades. Overall, our results emphasize that FEF inactivation decreases the overall count of saccade-related spikes across the population of iSC neurons, and in doing so, provides additional support for the fact that the iSC is putting out fewer spikes for metrically-matched saccades during FEF inactivation.

## Discussion

The saccadic spatiotemporal transformation is traditionally viewed as a function of the oculomotor brainstem, occurring between the iSC and the downstream burst generator (Groh, 2001; Moschovakis et al., 1998; Scudder et al., 2002). The linear dynamic ensemble-coding model proposed by Goossens and Van Opstal (2006) incorporates this view, relegating the role of cortical inputs like the FEF to specification of saccade goal, which is then executed by the iSC and burst generator without cortical involvement. We tested a core prediction of this model using reversible FEF inactivation: if the saccade displacement vector is the same, then the overall number of iSC spikes should be equivalent during FEF inactivation. Instead, we found that FEF inactivation reduced the number of iSC spikes for displacement-matched saccades, both at the center and throughout the entirety of the movement field, doing so in a task-dependent manner. Fundamentally, the iSC emits fewer spikes for displacement-matched saccades during FEF inactivation. In the following sections, we discuss the implications of our results from the perspective of the model, and from the perspective of how the oculomotor system may instantiate the spatiotemporal transformation.

### A task-dependent role for the FEF in the spatiotemporal transformation

Our four tasks varied the need to delay a response and remember the location of the peripheral target. We found that the decrease of saccade-related iSC spikes for metrically-matched saccades during FEF inactivation scaled with the cognitive demands of the task, with the greatest decrease accompanying memory-guided saccades that required a delayed response to a remembered target location and the smallest decrease accompanying direct saccades that required an immediate response to a persistently visible target. These task-dependent effects mirror the impact of FEF inactivation in more cognitively-demanding saccade tasks (Dias and Segraves, 1999; Peel et al., 2014; Sommer and Tehovnik, 1997). Further, we found that the spatial distribution of saccade-related iSC activity did not change during FEF inactivation. These results could not have been foreseen, as the decreases in iSC spike frequency during FEF inactivation that accompany lower saccade velocities (Peel et al., 2017) could have been offset by increases in saccade duration to yield a fixed number of spikes. Inputs to the iSC from extra-tectal sources also persist during FEF inactivation, but any compensatory changes in such inputs, in conjunction with intrinsic circuits within the iSC, are apparently insufficient to ensure a fixed number of iSC spikes during FEF inactivation.

Van Opstal and colleagues tested the linear dynamic ensemble coding model using a perturbation approach where a blink was induced just prior to a direct visually-guided saccade (Goossens and Van Opstal, 2000a). They found that the iSC continued to emit a fixed number of spikes despite remarkably perturbed saccadic trajectories (Goossens and Van Opstal, 2006). In contrast, we find that the number of iSC spikes changes during FEF inactivation, despite relatively modest decreases in saccade velocity and a straight saccade trajectory. We speculate that the differences in results arising from blink-perturbation versus FEF inactivation relate to the level at which each perturbation influences brain function. In contrast with direct manipulation of cortical activity via cryogenics, the trigeminal blink reflex appears to interact with iSC activity within less than 10 ms, presumably via subcortical trigiminotectal, cerebellotectal, or nigrotectal pathways, and also influences saccade trajectory via influences exerted downstream of the iSC (Goossens and Van Opstal, 2000b).

Previous studies on the linear dynamic ensemble coding model relied on saccades made directly to presented targets (Goossens and Van Opstal, 2006, 2012). Our warm-to-warm comparison, which established the noise inherent to our saccade matching procedure, shows that iSC neurons emits a fixed number of spikes for metrically-matched saccades in all paradigms, including memory and delayed visually-guided saccades. Unfortunately, the majority of our neurons were only tested in a single behavioural task, given our specific focus on FEF inactivation. Given the task-dependent nature of our results, a future test for the linear dynamic ensemble coding model will be to determine whether a given iSC neuron emits a fixed number of spikes throughout a response field for metrically-matched saccades generated in different paradigms with varying degrees of cognitive involvement (e.g., comparing direct versus delayed visually-guided saccades, or pro-versus anti-saccades). Doing so would clarify the contributions of extra-tectal sources to saccadic control across a variety of tasks.

### An increased feedforward gain may counteract decreased iSC activity for metrically-matched saccades

Our results lead us to speculate as to what modifications to the linear dynamic ensemble-coding model would be required to explain the effects of FEF inactivation. Within the model, each iSC spike is applied independently (see Equation 1 in **Methods**), so that the brainstem burst generator evokes a saccade commanded by the summed mini-vectors associated with each iSC spike. The model proposes that iSC activity is read out by the brainstem circuitry by a total of five parameters (scaling factors for horizontal and vertical component based on projection strength, feedforward gain of saccade burst neurons, and fixed delays of brainstem activation and of the feedback loops). The implication of fewer iSC spikes for metrically-matched saccades during FEF inactivation is that the impact of each spike would have to be, somewhat paradoxically, greater. Of the five parameters, the simplest explanation of our results is that FEF inactivation increases the feedforward gain of saccade burst neurons in the brainstem, doing so in a task-dependent manner. Although we cannot rule out the influence of FEF inactivation on the other four parameters, changes to these parameters seem less likely for a variety of reasons. The horizontal and vertical scaling factors are based on a nonlinear representation of visual space within the iSC (Robinson, 1972) that presumably relate to projection strengths with the brainstem (Moschovakis et al., 1998; Ottes et al., 1986); it seems unlikely that these projection strengths could change suddenly during FEF inactivation. Further, as we found that FEF inactivation reduced iSC spike counts within time windows of various lengths and shifts relative to saccade onset and offset, it is not immediately obvious how changes in delay parameters could counteract the decreased number of iSC spikes arriving at the brainstem burst generator.

### Neurophysiological implications for the spatiotemporal transformation

Although the spatiotemporal transformation for saccades is traditionally viewed as a function of the oculomotor brainstem (Groh, 2001; Moschovakis et al., 1998; Scudder et al., 2002), the work of Schiller and colleagues (1980) showed that monkeys with SC ablations generate accurate saccades after a period of recovery, so long as the FEF was intact. Given that the FEF, like the iSC, represents saccade targets in a spatial reference frame, these results demonstrate that the spatiotemporal transformation can occur between the FEF and oculomotor brainstem in the absence of the iSC, after a period of recovery. Much remains to be learned about the role of signalling conveyed along the FEF pathway that bypasses the iSC, although this pathway is not sufficient to evoke saccades in the intact animal (Hanes and Wurtz, 2001). Regardless, our results are inconsistent with a straightforward idea that FEF signals duplicate what is issued by the iSC. Indeed, had FEF spikes also provided mini-vectors that sum with those from the iSC, then one could have predicted that more, not fewer, iSC spikes would have been required during FEF inactivation for metrically-matched saccades. Instead, our results suggest that FEF and iSC signals to the brainstem circuitry have unique contributions to the spatiotemporal transformation in an intact animal. This perspective complements findings from previous studies suggesting that the FEF and iSC monosynaptically connect with different regions within the oculomotor brainstem: neurons from the caudal iSC excite saccadic burst neurons within the paramedian pontine reticular formation at monosynaptic latencies (Raybourn and Keller, 1977), whereas FEF corticopontine neurons terminate within the brainstem region containing the omni-pause neurons (OPNs) (Segraves, 1992).

There are multiple potential mechanisms by which FEF inactivation could influence the readout of iSC activity. While speculative, one explanation for our results could be that FEF inactivation decreases the tonic level of OPN activity during stable fixation. This speculation is based on our observations that FEF inactivation decreases all aspects of functionally-defined ipsilesional iSC emanating from both the rostral and caudal iSC (Dash et al., 2018; Peel et al., 2017), and ideas on how OPNs receive scaled excitatory inputs from the rostro-caudal extent of the iSC (Everling et al., 1998; Gandhi and Keller, 1997). Given the mutually-antagonistic relationship between OPN and burst neuron firing, such a decrease in OPN activity may lead to proportionate disinhibition of the brainstem burst generator, which could be the correlate of an increase in feedforward gain. It would also be very interesting to assess the impact of FEF inactivation on signaling conveyed to and within the cerebellum (Huerta et al., 1986; Xiong et al., 2002) given its widespread connections with the oculomotor brainstem and role in influencing saccade amplitude (Fuchs et al., 1993; Sato and Noda, 1992). Fortunately, recordings in both the OPNs and cerebellum during FEF inactivation are tractable, permitting new ways to test and refine biologically-plausible models for signal transformations within the oculomotor system.

## Acknowledgements

This work was supported by operating grants from the Canadian Institutes of Health Research to BDC (MOPs: 93796, 123247 and 142317) and the Natural Sciences and Engineering Research Council (NSERC; RGPIN-311680). TRP was supported by an Ontario Graduate Scholarship and TRP and SD were supported by funding from an NSERC CREATE grant.

